# Potent synthetic lethality between PLK1 and EYA-family inhibitors in tumours of the central and peripheral nervous system

**DOI:** 10.1101/2025.01.19.633804

**Authors:** Christopher B. Nelson, Jadon K. Wells, Ekaagra Kesarwani, Alexander P. Sobinoff, Jixuan Gao, Karen L. MacKenzie, Rebecca C. Poulos, Urwah Nawaz, Xiang Wang, Heide L. Ford, Hilda A. Pickett

## Abstract

The Eyes Absent family of protein phosphatases (EYA1-4) are aberrantly expressed and tumour-promoting across many devastating cancers of neurological origin affecting both children and adults. It has recently been demonstrated that EYA1 and EYA4 promote tumour cell survival by increasing the active pool of Polo-like kinase 1 (PLK1) molecules. This discovery provides a rationale for the therapeutic combination of EYA inhibitors with direct, ATP-competitive, PLK1 inhibitors. Here, we demonstrate potent and synergistic effects of EYA and PLK1 inhibition in cancer cell lines that overexpress EYA1 and/or EYA4, including in neuroblastoma and glioblastoma models. We identify decreases in PLK1 activity and RAD51 foci formation, and increases in mitotic arrest and cell death, as mechanistic contributors to combination sensitivity. Combined EYA and PLK1 inhibition is also effective in glioblastoma stem cell models that overexpress EYA1/EYA4 and specifically targets the cancer stem cell state. Finally, through multi-omic correlational analysis, we identify high levels of the NuRD complex and SOX9 as contributors to combination treatment sensitivity. Overall, this work identifies a novel synthetic lethal combination therapy with potential utility across a wide range of neurological cancers.

## Introduction

During development, EYA1-4 serve as integral components of the retinal determination gene network that contributes to cell-fate determination across various tissues^1, 2^. While expression of most of the EYAs is largely restricted in normal adult tissues, emerging evidence indicates that these proteins are frequently overexpressed and tumorigenic in cancers of both the peripheral and central nervous systems, including malignant peripheral nerve sheath tumours^3^, neuroblastoma^4^, low grade glioma/astrocytoma^5, 6^, glioblastoma^7–9^, and medulloblastoma^10–12^. Monotherapy with pan-EYA family phosphatase inhibitors has been tested in glioblastoma and medulloblastoma xenograft models, where EYA inhibition has been shown to increase survival^7,11^. However, targeted monotherapies almost always result in acquired resistance, and there is a growing appreciation for the need to identify rational combination therapies for effective cancer treatment^13–15^.

PLK1 is an essential mitotic kinase that functions during centrosome maturation, mitotic entry, checkpoint recovery, spindle formation, and other processes^16, 17^. PLK1 is also an important cancer target for many of the same tumours as the EYAs, and direct, ATP-competitive, PLK1 inhibitors are effective in preclinical models of glioblastoma^18–22^, neuroblastoma^23–26^, and medulloblastoma^27–30^. However, PLK1 inhibitor-based monotherapies have not been clinically successful, and there is an emerging shift towards the inclusion of PLK1 inhibitors in combination therapies^31, 32^. Recently, we found that PLK1 is a substrate of both EYA1 and EYA4^33^.

EYA-mediated dephosphorylation of PLK1 at tyrosine 445 modifies its polo-box domain and increases the interaction between PLK1 and PLK1 activation complexes^33^. This pathway increases the active pool of PLK1 molecules and stimulates mitotic progression in cancer cells that express EYA1 and/or EYA4^33^. Therefore, EYA-family inhibitors represent an indirect strategy for PLK1 inhibition in a subset of tumours. We reason that reducing the active pool of PLK1 molecules with EYA inhibitors may further sensitize cancer cells to direct PLK1 inhibitors and that this combination will specifically target cancers with EYA1 and/or EYA4 overexpression. EYA1 and EYA4 also share non-mitotic functions with PLK1, including the promotion of RAD51 filament formation during DNA repair, and the attenuation of apoptotic signalling^34^. Therefore, co-targeting of EYAs with PLK1 has multiple potential mechanisms of action.

In this study, we investigated whether the PLK1 inhibitors BI2536 and volasertib are synthetically lethal with the pan-EYA inhibitor benzarone in cancer cells expressing high levels of EYA1 and/or EYA4. We identified potent synthetic lethality in both a mixed cancer-type panel of cancer cell lines (including neuroblastoma and glioblastoma) and a panel of patient-derived glioblastoma stem cell lines. Key phenotypic effects of the combination treatment included decreased PLK1 substrate phosphorylation and reduced RAD51 accumulation on DNA. Ultimately, this led to an increase in mitotic arrest and apoptosis. In addition, we identified cancer stemness, SOX9, and the NuRD complex as important determinants of sensitivity to the combination treatment.

Overall, these data provide proof of principle for combined EYA and PLK1 inhibition, highlighting the potential utility of this therapeutic strategy for the treatment of glioblastoma, neuroblastoma, and other neurologically derived tumours characterized by overexpression of the EYA family.

## Results

### High expression levels of EYA1 and EYA4 confer sensitivity to combination EYA and PLK1 inhibition

To determine whether expression levels of EYA1 and EYA4 are predictive of sensitivity to the combination of pan-EYA and PLK1 inhibitors, we used publicly available transcriptomic data to assemble a panel of cancer cell lines with high EYA1 and EYA4 mRNA expression (“high expressors,” mean EYA1 and EYA4 expression > 2.5 log2 TPM +1, n=8) or low/undetectable EYA1 and EYA4 expression (“low expressors,” mean EYA1 and EYA4 expression < 0.25 log2 TPM +1, n=7) (Fig. 1A, expression data: https://depmap.org/portal/, 22Q4). To validate these groups, we performed western blots of all four EYAs across the 15 cell lines (Fig. 1B). Consistent with public transcriptomic data, the high expressor group had significantly higher EYA1/4 protein levels relative to the low expressor group (Fig. 1C, ** p ≤ 0.01,). Expression of EYA2 and EYA3 was relatively consistent across all cell lines and did not significantly differ between groups (Fig. 1B, EV1A-B).

**Figure 1:**
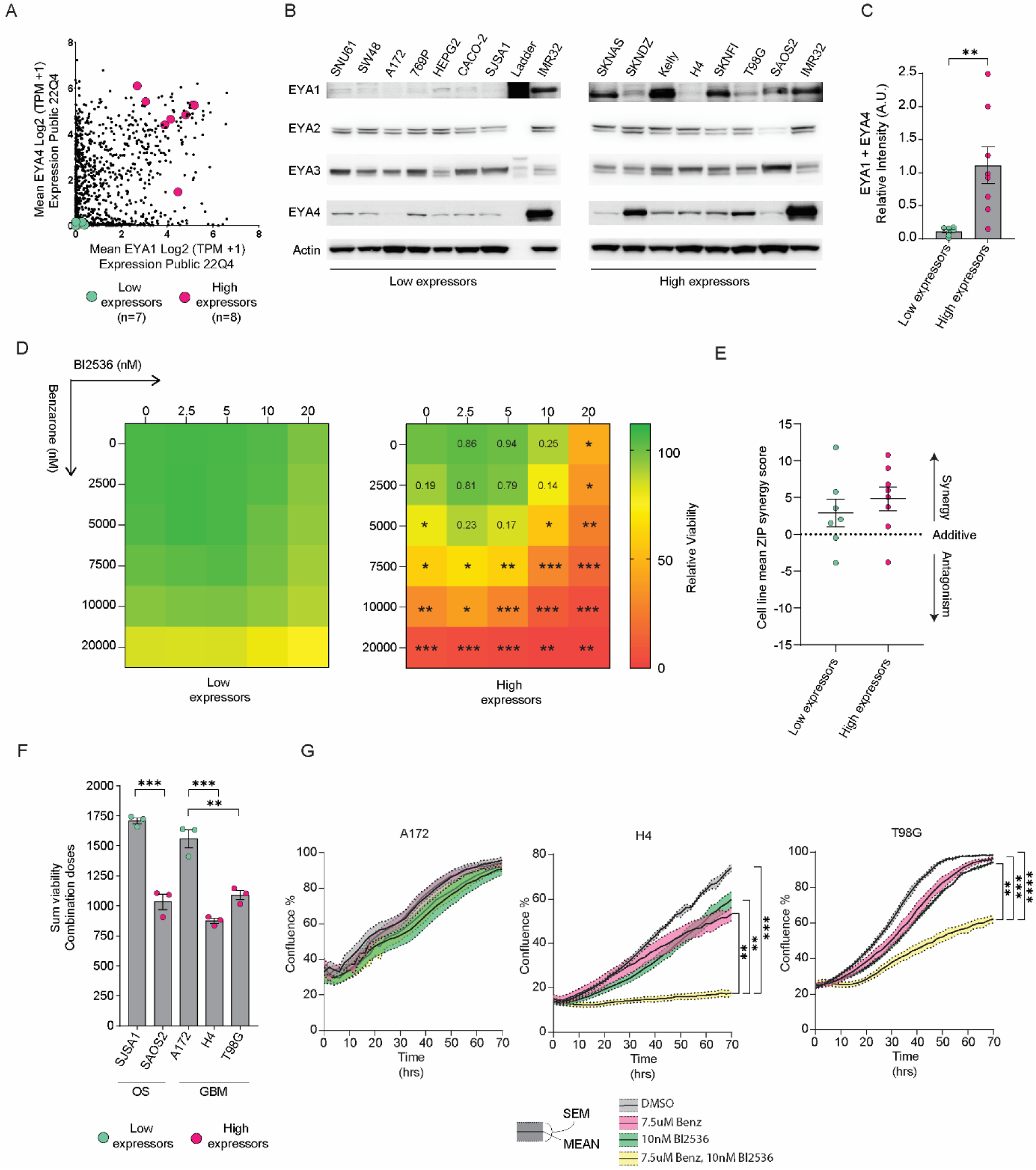
Combined EYA and PLK1 inhibition potently and synergistically reduces viability of cancer cell lines with high EYA1 and EYA4 expression. **(A)** Cell lines identified as having low expression of EYA1 and EYA4 (“low expressors,” n= 7, pink) or high expression of EYA1 and EYA4 (“high expressors,” n=8, green) were identified from public RNA-seq data of 1408 total cell lines. **(B)** Western blots of EYA1-4 across all low and high expressor cell lines. **(C)** Quantification of sum EYA1 and EYA4 proteins levels relative to IMR32 via densitometry (two-tailed *t* test, ** p ≤ 0.01). **(D)** Relative viability across combination dose-response matrixes measured by alamarBlue (two-tailed *t* tests were used to compare viability at each dose, * p ≤ 0.05, ** p ≤ 0.01, *** p ≤ 0.001). **(E)** Mean ZIP synergy score of all low or high expressor cell lines. **(F)** Sum viability across all combination doses in two cancer type-matched cell line sets including osteosarcoma (OS) and glioblastoma (GBM) (two-tailed *t* tests were used to compare sum viability between each low expressor cell line and cancer type-matched high expressors, ** p ≤ 0.01, *** p ≤ 0.001). **(G)** Quantitation of mean cell confluence over time following treatment of GBM cell lines with DMSO, Benzarone (Benz), Bi2536, or Benzarone and Bi2536, at the indicated doses (n=3, coloured regions indicate standard errors of the mean, one-way ANOVA was used to compare the area under the curve between the combination treatment, individual drug treatments and DMSO, ** p ≤ 0.01, *** p ≤ 0.001, **** p ≤ 0.0001).

Evidence for binding of the pan-EYA inhibitor benzarone to the EYA proteins has previously been limited to *in-vitro* assays and indirect phenotypic effects^33, 35, 36^. Here, we employed the cellular thermal shift assay (CETSA), to directly demonstrate binding of benzarone to each of the EYAs in IMR32 cells, indicative of on-target activity (EV1C-D).

Each cell line in our panel was treated with a benzarone and BI2536 combination dose-response matrix and cell viability was examined using the alamarBlue assay. EYA1/4 high expressor cell lines were significantly more sensitive to increasing doses of either benzarone or BI2536 alone, as well as to increasing combination doses of the two drugs, when compared to low expressors (Fig. 1D, * ≤ 0.05, ** ≤ 0.01 *** ≤ 0.001). We then computed the zero interaction potency (ZIP) synergy score for all cell lines. While the high expressor cell lines trended towards higher mean ZIP synergy scores, both groups had a positive mean ZIP synergy score and no statistically significant difference between groups was observed (Fig 1E). These data indicate that the drug combination was broadly synergistic, but was effective at lower doses in cell lines with high levels of EYA1 and/or EYA4. To determine if the synergistic effects of this drug combination were specifically related to treatment with BI2536, we performed dose-response matrices and viability testing in H4 and T98G cell lines using volasertib as an alternative PLK1 inhibitor. Benzarone and volasertib was also found to be a synergistic combination with mean synergy scores of 12.14 and 5.60 in the respective cell lines (EV1E-F).

To account for potential tumour type-specific effects, matched sets of cell lines with low or high EYA1/4 expression were included for both osteosarcoma (OS) and glioblastoma (GBM). We compared the sum viability across all combination doses within individual cell lines of the matched sets. In both OS and GBM, high expressor cell lines (SAOS2, H4 and T98G) had significantly lower sum viability compared to matched low expressor cell lines (SJSA1, A172, Fig. 1F, ** p ≤ 0.01, *** p ≤ 0.001). To further investigate the effects of the drug combination in the GBM cell lines, we performed live cell imaging following treatment with a single concentration of DMSO, benzarone, BI2536 or a combination of the drugs. The individual drug concentrations for this experiment (benzarone 7.5 µM, BI2536 10 nM) caused a small but detectible effect on cell viability in the GBM cell lines (means of 10.28% and 6.60%, respectively) and were chosen to maximize the potential for detecting a combinatorial effect on cell growth. Consistent with the observed effects on viability, the combination treatment significantly reduced cell growth compared to either individual drug or DMSO control in H4 or T98G high expressor GBM cell lines, but not in A172 low expressor cells (Fig. 1G, ** p ≤ 0.01, *** p ≤ 0.001, **** p ≤ 0.0001). Taking these results together, we conclude that combined inhibition of EYAs and PLK1 is potent and synergistic in cancer cell lines that overexpress EYA1 and/or EYA4.

### Combination EYA and PLK1 inhibition results in mitotic arrest, apoptosis, and genomic instability in GBM cell lines

To investigate how benzarone and BI2536 synergize to reduce cell growth and viability, we explored effects on the cell cycle using T98G cells expressing the FUCCI(CA)2 cell cycle indicator system across the combination dose-response matrix. DMSO treatment caused an increase in G1 cells over time, an expected outcome of contact inhibition accompanying increasing cellular confluence (Fig. 2A, n=4). Consistent with a mitotic arrest phenotype, increasing doses of benzarone, BI2536 and combination treatments produced immediate increases in the G2/M cell proportion which initially peaked at around 16 hours post-treatment (Fig. 2A, n=4). At higher drug concentrations, the 16-hour peak in G2/M cells was followed by a transient reduction, and then a progressive increase for the duration of the experiment (Fig. 2A, n=4). The percentage of G2/M cells at the initial peak (16 hrs) and at the final timepoint (72 hrs) were consistently greater in combination treatments compared to commensurate individual drug treatments, indicating that combination treatments trigger a more severe mitotic arrest phenotype (Fig 2A-B, n=4). To determine whether the combination treatment was cytotoxic, we examined Annexin V staining as a marker of apoptosis over time and across the dose-response matrix. The percentage of Annexin V positive cells increased with escalating doses of either individual drugs or combination treatments, following a roughly sigmoidal curve over time (Fig 2C, n=4). A more striking increase in the percentage of Annexin V positive cells was observed in most of the combination treatments compared to the commensurate individual doses, although this was not observed at the highest doses of benzarone (Fig 2C-D, n=4).

**Figure 2:**
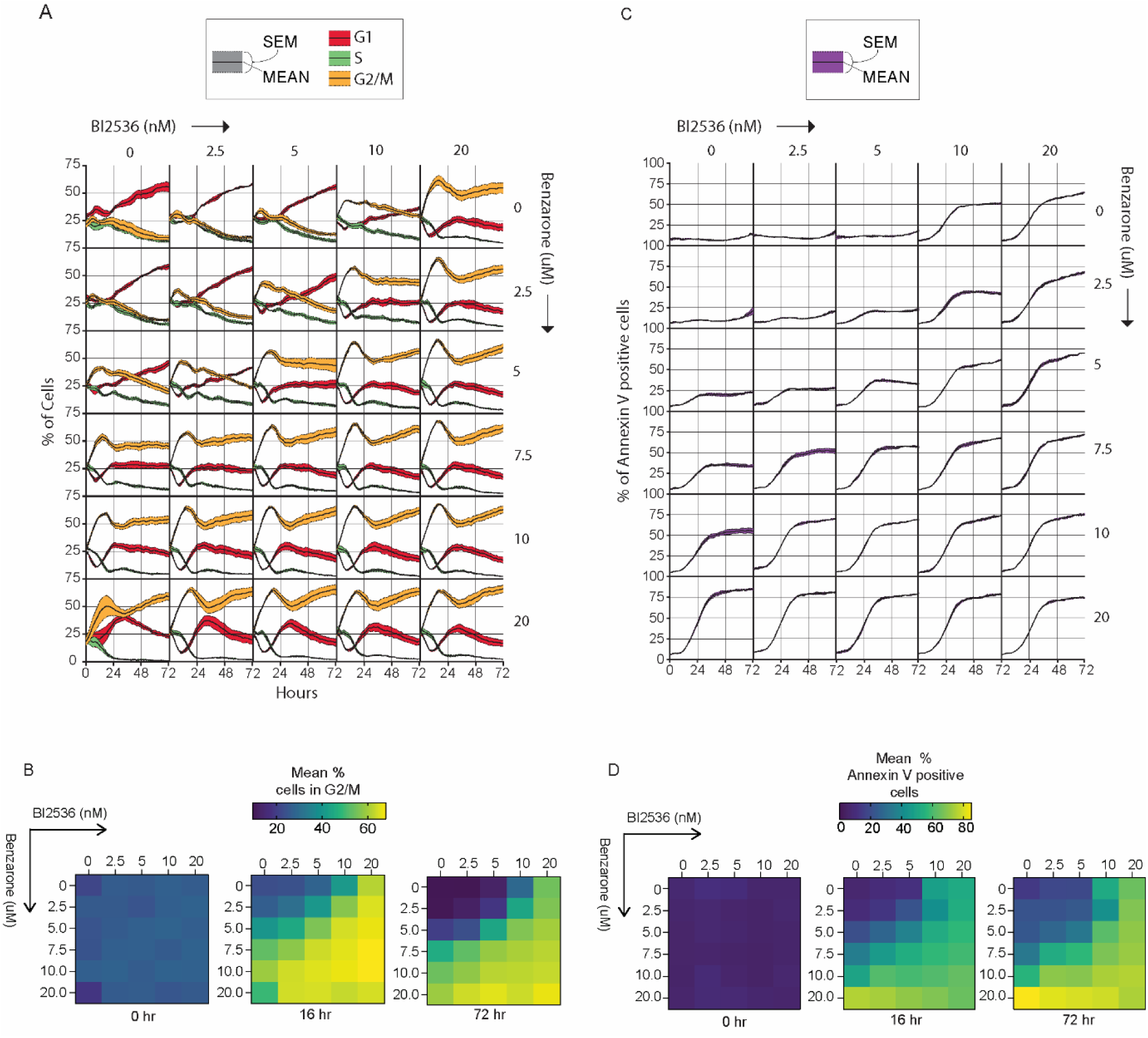
Combined EYA and PLK1 inhibition increases mitotic cell fraction and cell death in a dose and combination-dose dependent manner. **(A)** Mean percentage of T98G cells expressing the FUCCI(CA)2 cell cycle indicator showing the percentage of cells in G1, S, and G2/M phase over 72 hrs following treatment with benzarone, Bi2536, or combinations of both drugs (black lines with standard errors color-coded for each cell cycle phase). **(B)** Mean percentage of cells in G2/M phase across the dose-response matrix at 0 hrs, 16 hr, and 72 hrs following treatment. **(C)** Mean percentage of Annexin V positive T98G cells over 72 hrs following treatment with benzarone, Bi2536, or combinations of both drugs (black lines with standard error shown in purple). **(D)** Mean percentage of Annexin V positive cells across the dose-response matrix at 0 hrs, 16 hr, and 72 hrs following treatment.

As the observed G2/M arrest and apoptotic response is consistent with combinatorial reductions in PLK1 activity, we next examined the production of micronuclei, a form of genomic instability that can result from PLK1 inhibition^37–39^. Since T98G cells are hyper-pentaploid and have highly pleiomorphic nuclei, we investigated micronuclei in H4 cells, which have a consistent nuclear morphology that enables more reliable scoring. While 24-hour treatment with benzarone alone significantly increased micronuclei formation in H4 cells, the combination of benzarone and BI2536 caused a further increase in micronuclei, which was statistically significant compared to either drug treatment alone or DMSO control (EV2A-B *p ≤ 0.05, ** p ≤ 0.01, *** p ≤ 0.001, **** p ≤ 0.0001). Evidence of genomic instability was also observed by western blot analysis of H4 cells, in which the DNA damage marker gamma-H2AX was strikingly elevated in drug combination treated cells (EV2C). H4 cells treated with the drug combination also had increased levels of cleaved PARP and S10-phosphorylated H3, markers of cell death and mitotic arrest, respectively (EV2C).

### Combination EYA and PLK1 inhibition impacts PLK1 substrate phosphorylation and RAD51 foci formation in GBM cells

To determine whether combined treatment with benzarone and BI2536 additively disrupts PLK1 kinase activity in G2 and M-phase cells, we used a quantitative image-based cytometry approach in which G2 and M-phase cells were gated based on the staining intensity of DAPI and S10 phosphorylation of H3 in H4 cells. In G2 cells treated with inhibitors for a short duration (5.5 hrs), p-S46 TCTP, a model substrate of PLK1, was reduced following treatment with benzarone and further reduced in cells treated with the combination of benzarone and BI2536 (Fig. 3A-C, ** p ≤ 0.01, **** p ≤ 0.0001)^40–43^. In M-phase cells, p-S46 TCTP staining was reduced specifically in cells treated with the drug combination (Fig. 3D, * p ≤ 0.05). These results suggest that the combination treatment additively reduced PLK1 activity. BI2536 treatment alone did not reduce p-S46 TCTP in either G2 or M-phase cells, but significantly increased p-S46 TCTP relative to DMSO in M-phase (Fig. 3C-D, ** p ≤ 0.01). While p-S46 TCTP is a well-validated PLK1 substrate, it has not been examined following treatment with 10 nM BI2536, with previous reports indicating that 100 nM is sufficient for its reduction ^40–43^. As an additional control, we confirmed that treatment with 100 nM BI2536 could reduce mitotic p-S46 staining in H4 cells (EV3A, **** p ≤ 0.0001).

**Figure 3:**
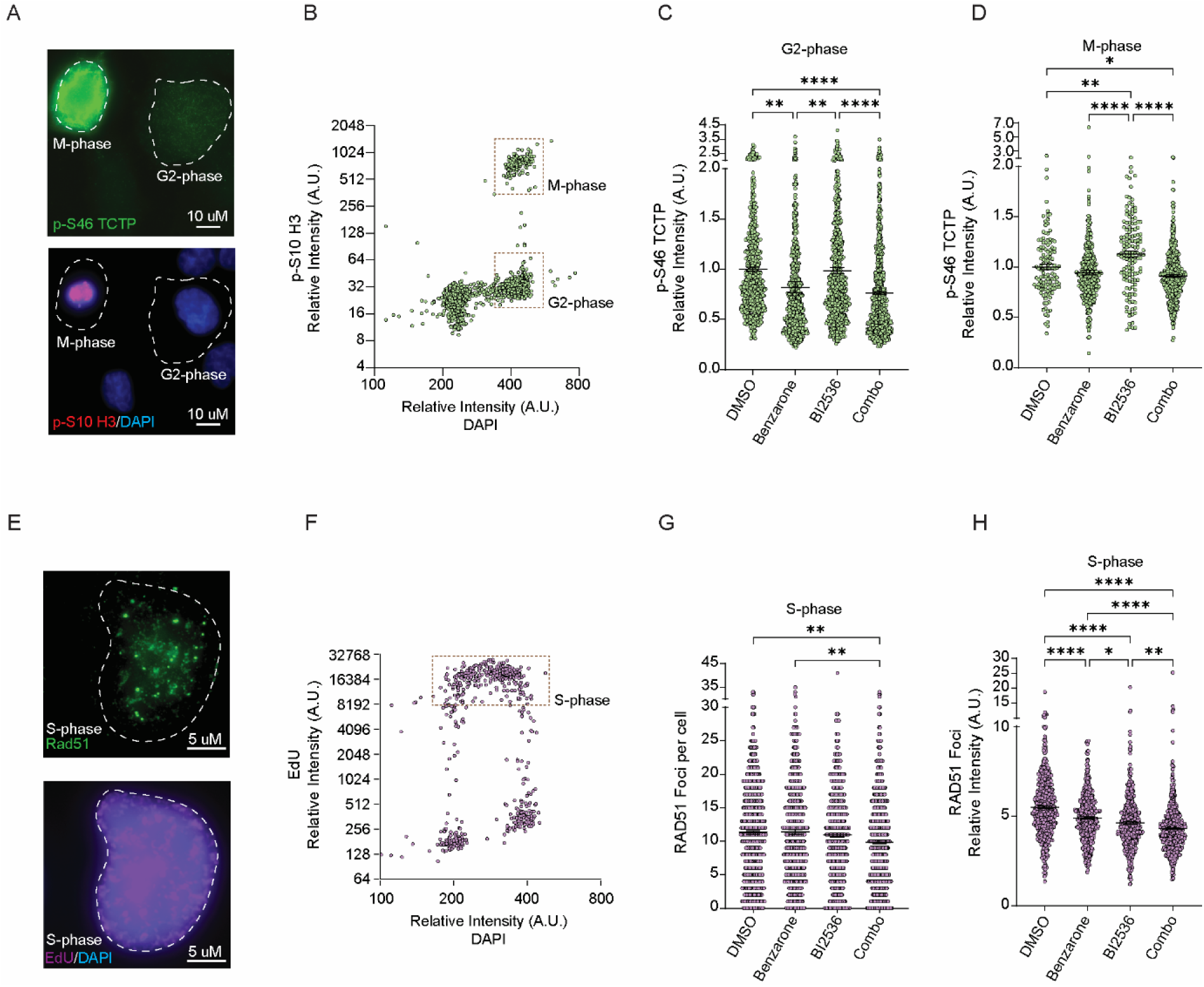
Combined EYA and PLK1 inhibition additively decreases PLK1 substrate phosphorylation in G2 and M-phase and RAD51 foci formation and intensity in S-phase. **(A)** Representative immunofluorescence images showing G2 and M-phase cells stained with p-S46 TCTP, p-S10 H3 and DAPI. **(B)** Representative quantitative image-based cytometry plot showing G2 and M-phase cell cycle gates as determined by staining intensity of p-S10 H3 and DAPI. **(C)** Cellular quantitation of p-S46 TCTP intensity in G2-phase following treatment with DMSO, Benzarone (7.5 µM), Bi2536 (10 nM), or Benzarone + Bi2536 for 5.5 hrs (one-way ANOVA with Tukey’s multiple comparison test, ** p ≤ 0.01, **** p ≤ 0.0001). **(D)** Cellular quantitation of p-S46 TCTP intensity in M-phase following treatment with DMSO, Benzarone (7.5 µM), Bi2536 (10 nM), or Benzarone + Bi2536 for 5.5 hrs (one-way ANOVA with Tukey’s multiple comparison test, * p ≤ 0.05, ** p ≤ 0.01, **** p ≤ 0.0001). **(E)** Representative immunofluorescence images showing an S-phase cell stained with RAD51, EdU, and DAPI. Individual RAD51 foci are indicated with red outline. **(F)** Representative quantitative image-based cytometry plot showing S-phase cell cycle gate as determined by staining intensity of EdU and DAPI. **(G)** Cellular quantitation of RAD51 foci number in S-phase following treatment with DMSO, Benzarone (7.5 µM), Bi2536 (10 nM), or Benzarone + Bi2536 for 5.5 hrs (one-way ANOVA with Tukey’s multiple comparison test, ** p ≤ 0.01). **(H)** Quantitation of RAD51 foci intensity in S-phase following treatment with DMSO, Benzarone (7.5 µM), Bi2536 (10 nM), or Benzarone + Bi2536 for 5.5 hrs (one-way ANOVA with Tukey’s multiple comparison test, * p ≤ 0.05, ** p ≤ 0.01, **** p ≤ 0.0001).

Both PLK1 and EYA4 promote RAD51 foci formation during homologous recombination (HR), a process that occurs in S or G2-phase cells and supports cancer cell growth and survival^44–48^. Using DAPI intensity and pulse incorporated EdU, we gated cells into S and G2 populations and analysed RAD51 foci. Following 5.5 hr inhibitor treatments, S-phase cells had significantly fewer RAD51 foci in combination treated cells relative to DMSO or benzarone alone (Fig. 3E-G, ** p ≤ 0.01). Additionally, the intensity of individual RAD51 foci was significantly reduced in S-phase cells treated with the drug combination compared to DMSO, benzarone or BI2536 alone (Fig. 3H, * p ≤ 0.05, ** p ≤ 0.01, **** p ≤ 0.0001). In G2 phase cells, the results trended in the same direction with a significant reduction in RAD51 foci intensity in the combination treated cells, however RAD51 foci number was not significantly reduced relative to DMSO and was only reduced relative to benzarone alone (EV3B-D, * p ≤ 0.05, ** p ≤ 0.01).

To determine if the observed reductions in RAD51 foci in drug combination treated cells was impacting the frequency of HR events, we evaluated HR-dependent DNA crossovers by quantifying sister chromatid exchange (SCE) events. Surprisingly, benzarone treatment significantly elevated SCEs, with a further increase in SCEs observed in the combination treatment (EV3E-F, ** p ≤ 0.01, **** p ≤ 0.0001). Therefore, the observed reductions in RAD51 foci and foci intensity do not reflect a reduction in HR and may instead be indicative of impaired RAD51 functions at replication forks where RAD51 foci can also be indicative of its functions in replication fork protection and reversal^49–52^. Overall, these data demonstrate that reductions in PLK1 activity and RAD51 foci contribute to the cellular effects of the combination treatment.

### EYA and PLK1 inhibition are potent and synergistic in a subset of GBM stem cell models

GBM stem cells are at the apex of the GBM tumour cell hierarchy and are responsible for driving tumour formation and therapy resistance^53, 54^. Due to the effectiveness of combined EYA and PLK1 inhibition in GBM cell lines, we next tested our drug combination in a panel of 12 patient-derived GBM stem cell models^55, 56^. These models mimic the clinical diversity of GBM as they span GBM molecular subtypes, including four representatives each of proneural, classical, and mesenchymal tumours^55^. While overexpression of EYA2 in GBM stem cells has been previously reported, we found that the majority of GBM stem cell models also express high levels of EYA1 and EYA4, but not EYA3, relative to a non-stem GBM cell line (Fig. 4A-B)^7^. Across all 12 models, cell growth was significantly reduced by combined EYA and PLK1 inhibition relative to DMSO and was reduced more than the additive effect of single drug treatments in all but one model (Fig. 4C-E, * p ≤ 0.01, ** p ≤ 0.01, **** p ≤ 0.0001). The magnitude of the growth reduction following combination treatment was highly variable and models were loosely grouped as “slightly sensitive,” (>70% relative growth, 2/12 models), “moderately sensitive,” (30-70% relative growth, 5/12 models), or “highly sensitive,” (<30% relative growth, 5/12 models) (Fig. 4C-E). The highly sensitive group had representation from all GBM subtypes, but predominantly comprised classical subtype models, with 3/4 of the classical subtype models being highly sensitive to the combination treatment (Fig. 4C-E).

**Figure 4:**
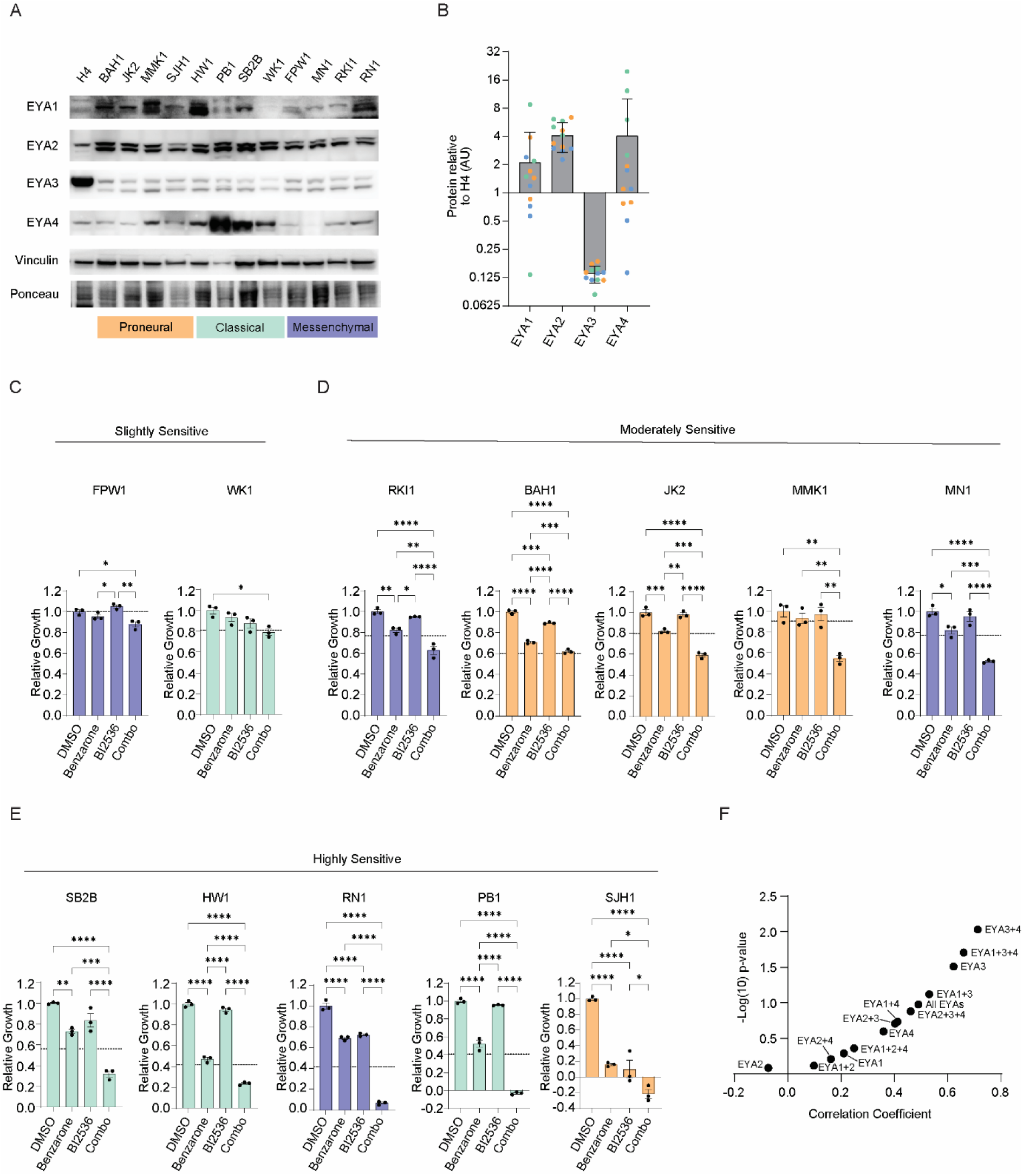
Patient-derived glioblastoma stem cell lines are sensitive to combined EYA and PLK1 inhibition. **(A)** Western blot of GBM stem cell models and H4 GBM cell line using the indicated antibodies. **(B)** Quantification of mean EYA1-4 proteins levels relative to H4 cells via densitometry. **(C-E)** Quantitation of relative growth over 4-5 days for 12 GBM stem cell models following treatment with DMSO, Benzarone (7.5 µM), Bi2536 (10 nM), or Benzarone + Bi2536 (one-way ANOVAs with Tukey’s multiple comparison test, * p ≤ 0.05, **p ≤ 0.01, *** p ≤ 0.001, **** p ≤ 0.0001). Dashed lines indicate the predicted additive effect (sum viability reduction of benzarone and BI2536 treatments). Models are grouped according to sensitivity to the combination treatment (combination relative growth >70% = “Slightly Sensitive,” 35-70% = “Moderately Sensitive,” ≤35% = “Highly Sensitive”). **(F)** Pearson correlation coefficients and -Log(10) p-values between viability loss and EYA protein Z-scores in GBM stem cell models following combination treatment.

Correlation analysis between the sensitivity to the combination treatment and the protein level of each EYA revealed positive associations with EYA1, 3, and 4, but not EYA2 (Fig. 4F). Surprisingly, EYA3 levels were the most highly correlated with combination sensitivity (Fig. 4F, r = 0.62, p = 0.03). Additionally, the sum Z-score of EYA1 and EYA4 had a higher correlation coefficient than either protein alone and Z-score combinations of EYA1, 3 and 4, and EYA3 and 4 were both more highly correlated with sensitivity than any individual EYA (Fig. 4F, EYA1 + 4: r = 0.41, p =0.18, EYA1+3+4: r = 0.66, p = 0.02, EYA 3+4: r = 0.71, p = 0.009).

### EYA and PLK1 inhibition targets the GBM stem cell state and causes tumour spheroid shrinkage

To determine whether the combination treatment specifically targeted the GBM stem cell state, we differentiated the GBM stem cell model PB1 by removal of supplemental growth factors (FGF, EGF) for 48 hrs. Differentiated PB1 cells were able to attach and grow on cell culture dishes without the application of matrigel and had reduced expression of GBM stem cell markers including CD133 and SOX9 (EV Fig 5A-B). PB1 stem cells were significantly more sensitive than differentiated PB1 cells to benzarone, BI2536 and combination drug treatments across a dose-response matrix (Fig. 5A, * p ≤ 0.05, ** p ≤ 0.01, *** p ≤ 0.001, **** p ≤ 0.0001, n=4). Furthermore, the mean combination synergy score was synergistic and significantly higher in PB1 stem cells compared to differentiated PB1 cells, for which the combination was found to be additive (Fig 5B, ** p ≤ 0.01, n=4).

**Figure 5:**
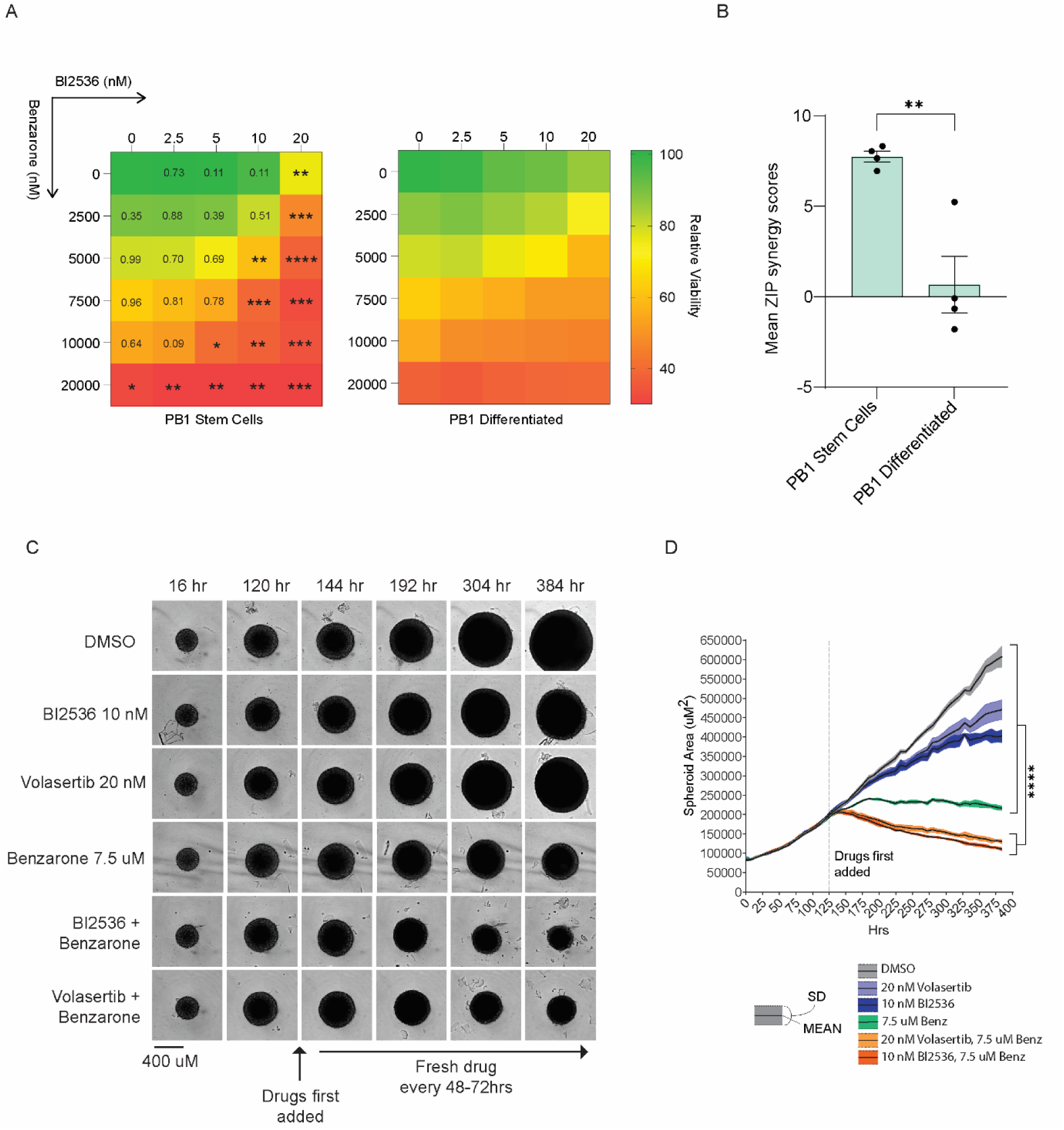
EYA and PLK1 inhibitors have enhanced potency and synergy towards the GBM stem cell state and cause tumour spheroid shrinkage. **(A)** Relative viability across combination dose-response matrixes measured by alamarBlue in PB1 GBM stem cells and PB1 GBM differentiated cells. (two-tailed *t* test were used to compare each dose, * p ≤ 0.05, ** p ≤ 0.01, *** p ≤ 0.001, **** p ≤ 0.0001). **(B)** Mean ZIP synergy scores of PB1 stem and differentiated cells (two-tailed *t* test, ** p ≤ 0.01). **(C)** Representative images of PB1 tumour spheroids. **(D)** Mean spheroid area over time across at least 3 spheroids per treatment (ANOVA with Šídák’s multiple comparisons test was performed on the final spheroid areas, **** p ≤ 0.0001).

GBM stem cells grown in round bottom plates form tumour spheroids that recapitulate *in-vivo* tumour growth. We next evaluated EYA and PLK1 inhibition in GBM stem cell-derived tumour spheroids. Individual inhibition of PLK1 or EYA reduced the rate of growth or caused a plateau in growth, respectively (Fig. 5C-D). In contrast, the combination of benzarone with either BI2536 or volasertib caused shrinkage of PB1-derived spheroids resulting in reduced spheroid size relative to all other treatments (Fig. 5C-D, **** p ≤ 0.0001). Together, these results demonstrate that the synergistic effects of EYA and PLK1 combination therapy in GBM stem cells are sufficient to shrink GBM tumour spheroids.

### Sensitivity to combination EYA and PLK1 inhibition is associated with expression of the NuRD nucleosome remodelling complex

EYA family member expression is unlikely to completely explain the observed differences in sensitivity to the combination treatment, as even some highly sensitive GBM stem cell models had moderate EYA levels (e.g., SJH1). Therefore, we interrogated published proteomic data available for the GBM models to identify other relevant proteins and pathways^57^. Correlations between GBM stem cell proteomes and sensitivity to EYA and PLK1 combination therapy were used to inform pathway and gene ontology analyses. Positively correlated proteins were highly enriched for chromatin remodelling complexes and pathways (Fig. 6A-C, enrichment score > 4). Many of these proteins are components of the nucleosome remodelling and deacetylase (NuRD) complex, of which all members within the proteomic dataset had a positive correlation with sensitivity (Figure 6A-D). To determine if this relationship is also detectable at the RNA level, we examined the correlation of NuRD complex members within three published microarray datasets^55^. This included data from the original tumours from which the GBM stem cell models were derived, data from the GBM stem cell lines grown in 2D culture, and data from xenografts of GBM stem cell lines^55^. Additionally, we examined RNA-seq data for the GBM stem cell lines (QCell resource: https://www.qimrberghofer.edu.au/commercial-collaborations/partner-with-us/qcell/)^55^. The magnitude of the mean NuRD component correlation with sensitivity was not as large at the mRNA level, however there was a positive mean correlation coefficient between NuRD components and combination sensitivity in all datasets, indicating a robust relationship in GBM (Fig. 6E).

**Figure 6:**
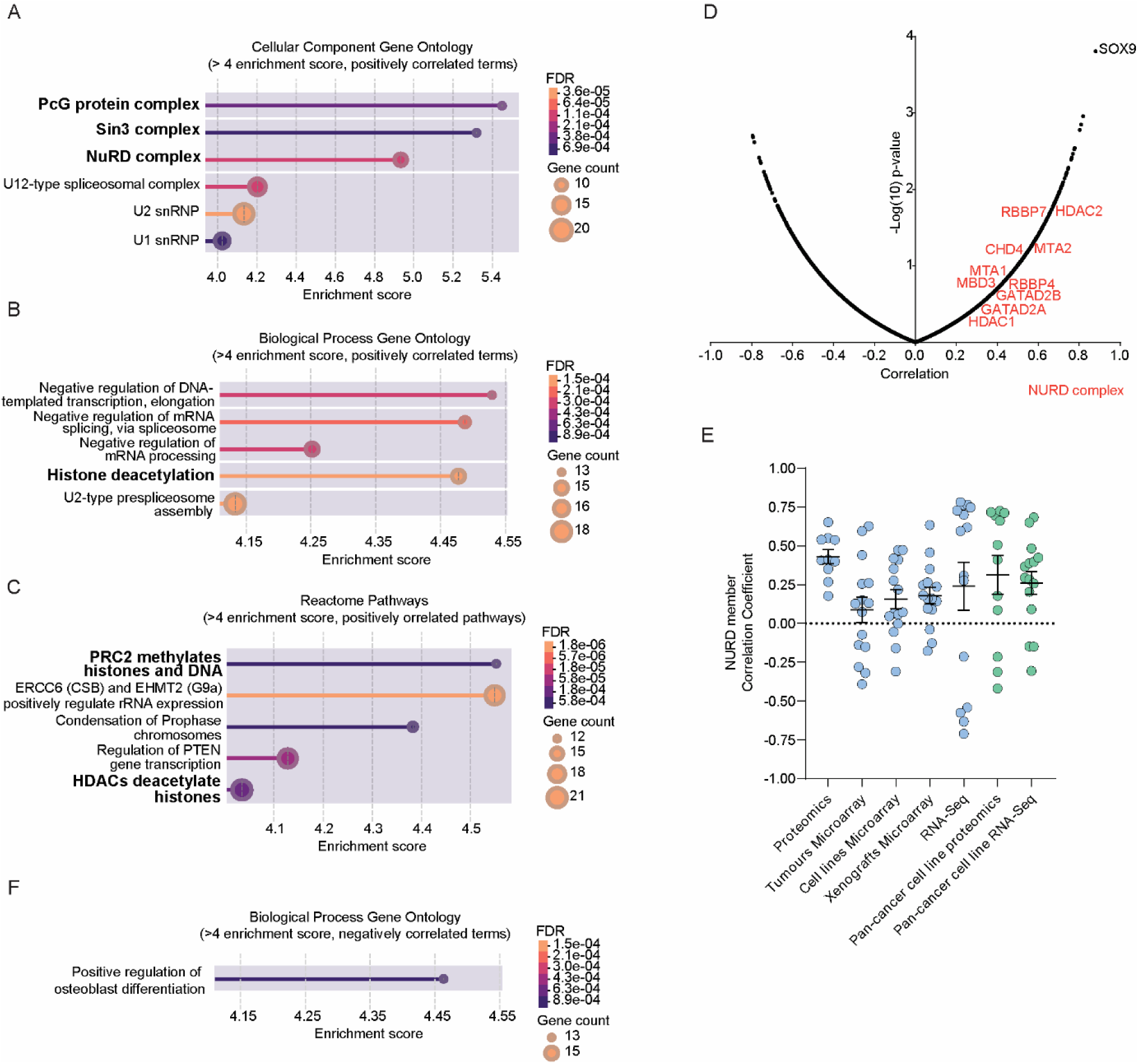
Gene-ontology and pathway associations with combination therapy responses. **(A)** Highly enriched cellular component gene ontology terms generated using the STRING proteins with values/ranks function. Terms related to proteins that were positively correlated with sensitivity to the combination therapy are presented (FDR ≤ 0.01, Enrichment score > 4, correlation coefficients between each protein and sensitivity to the combination was used as the value/rank term). **(B)** Highly enriched biological process gene ontology terms generated using the STRING proteins with values/ranks function. Terms related to proteins that were positively correlated with sensitivity to the combination therapy are presented (FDR ≤ 0.01, Enrichment score > 4, correlation coefficients between each protein and sensitivity to the combination was used as the value/rank term). **(C)** Highly enriched reactome pathways generated using the STRING proteins with values/ranks function. Pathways related to proteins that were positively correlated with sensitivity to the combination therapy are presented (FDR ≤ 0.01, Enrichment score > 4, correlation coefficients between each protein and sensitivity to the combination was used as the value/rank term). **(A-C)** Terms related to chromatin remodelling complexes are shown in bold. **(D)** Pearson’s correlation coefficients from regression analysis of individual proteins and sensitivity to the combination treatment and -Log(10) p-values. The most highly correlated individual protein, SOX9, is indicated in black, and components of the NuRD family are indicated in red. **(E)** NuRD complex member Pearson correlation coefficients with combination sensitivity from proteomic or transcriptomic datasets of 12 GBM stem cell models (blue) or from the 15 cell lines tested in Figure 1A-E (green) (mean correlation coefficient and SEM are presented, dots represent individual NuRD components). **(F)** Highly enriched biological process gene ontology terms generated using the STRING proteins with values/ranks function. Terms related to proteins that were negatively correlated with sensitivity to the combination therapy are presented. (FDR ≤ 0.01, Enrichment score > 4, correlation coefficients between each protein and sensitivity to the combination was used as the value/rank term).

To determine if the positive correlation between NuRD and combination sensitivity was a GBM-specific phenomenon or more broadly applicable across cancers, we examined NuRD component correlations in publicly available proteomic and transcriptomic data corresponding to the pan-cancer cell line panel described in Figure 1. In both datasets, we identified that the mean correlation coefficient determined from regression analysis of NuRD components and combination drug sensitivity was positive (Fig. 6E). Comparison of individual NuRD component correlations indicated that several individual NuRD members, including MTA1, MTA2, CHD4, GATAD2B, and MBD3 were positively correlated across all 7 datasets (EV Fig. 6A).

The only term or pathway with an enrichment score of > 4 for negatively correlated proteins was “positive regulation of osteoblast differentiation” (Fig. 6F). This may suggest that more differentiated and less “stem-like” GBM models are less sensitive to the combination treatment. This rationale is supported by the most highly correlated individual protein, SOX9, being a transcription factor that promotes GBM stemness (Figure 6D)^58–61^. Taken together these results highlight important pathways and proteins, including the NuRD complex and SOX9, that contribute to sensitivity to the combination treatment and may be useful for identifying sensitive tumours in the future.

## Discussion

Tumours of the central and peripheral nervous systems including GBM and NB are lethal malignancies with a scarcity of effective treatment options. PLK1 inhibition has repeatedly been shown to be a successful strategy in preclinical models of these tumours; however, PLK1 is essential to all dividing cells and there is a need to design rational combination treatments with PLK1 inhibitors to improve their potency in cancer cells^31, 32^. In this study, we describe the novel combination of PLK1 and EYA phosphatase inhibition. This combination strategy is both potent and synergistic in cancer models that overexpress EYA1 and/or EYA4, including NB and GBM cell lines, as well as patient-derived GBM stem cell models and tumour spheroids. We also find that the combination of PLK1 and EYA inhibitors specifically targets the GBM stem cell state, a highly sought-after therapeutic attribute that current clinical options do not effectively address.

This work builds on our previous discovery that EYA1 and EYA4 promote the activation of PLK1^33^. Consistent with a mechanism of action caused by combined direct and indirect inhibition of PLK1, we observed enhanced G2/M arrest and cell death as well as reduced PLK1 substrate phosphorylation and RAD51 filament formation. Interestingly, we found that a low dose of PLK1 inhibitor was not sufficient to reduce PLK1 substrate phosphorylation by itself, and instead increased p-S46 TCTP in mitotic cells. We reason that this may be due to adaptive dysregulation of PLK1 activation mechanisms in response to partial PLK1 inhibition. In this scenario the addition of an EYA inhibitor may help to overcome adaptation to PLK1 inhibitors, a plausible explanation given the role of the EYAs in promoting the PLK1 activation pathway.

Prior work suggests that both PLK1 and EYA4 activate RAD51 which promotes HR^44–48^. While combined inhibition of PLK1 and EYAs disrupted RAD51 foci formation, we found that it promoted HR, as indicated by SCE formation. This could be explained by a primary function of the EYAs and PLK1 in promoting RAD51 at replication forks rather than at double strand breaks. A deficit in RAD51 at replication forks could cause excessive replication fork breakage which may stimulate HR overall, even if individual HR events have a lower probability of success. This idea is supported by our observation that combination treated cells have highly elevated gamma-H2AX, a marker of double strand breaks.

As the potential utility of the combination treatment depends on the specific overexpression of EYA1 and/or EYA4 in cancer cells, we validated the relationship between combination treatment sensitivity and EYA1/4 expression across cancer cell lines and in GBM stem cell models. To our surprise the level of EYA3 was also associated with sensitivity to the combination treatment in GBM stem cells, despite having relatively low abundance across the models. It was recently shown that all four EYAs can interact with one another including formation of a heterotetrametric complex^62^. Therefore, EYA3 may function with EYA1 and EYA4 to promote PLK1 activation in an EYA-complex dependent manner. In support of this possibility, several combinations of EYAs were more strongly associated with sensitivity to the combination treatment compared to any individual EYA in GBM stem cell models.

While the levels of the EYAs are clearly important in predicting sensitivity to the combination treatment, some individual GBM models and cell lines with relatively modest levels of the EYAs were still sensitive. This is best exemplified by SJH1, which has average expression of the EYAs relative to other GBM stem cell models, but showed the greatest reduction in cell growth following the combination treatment. This indicates that other factors contribute to combination treatment sensitivity. We identified a strong association between high protein and RNA expression of the NuRD chromatin remodelling complex and combination treatment sensitivity in both the GBM stem cell models and the pan-cancer cell line panel. This suggests that combined inclusion of EYA and NuRD complex member levels could help to guide future use of the combination therapy irrespective of tumour type.

We also observed that GBM stem cell models of the classical transcriptional subtype were particularly sensitive to the combination treatment (3/5 highly sensitive models). Only 3 out of 15 NuRD components and none of EYA1, 3, or 4 are among the 840 genes commonly used for GBM subtype classification, which may suggest that the classical subtype has an independent relationship with combination sensitivity^63^. However, it is possible that expression levels of NuRD and EYA family members have subtle relationships to the classical subtype, or relationships that are masked by gene-family level redundancies.

Additionally, our results indicate that GBM stem cells are more sensitive to the combination treatment than their differentiated counterparts. These results suggest that GBM stem cells have a greater reliance on PLK1 activity and/or the EYA-mediated PLK1 activation pathway. Evidence that the combination targets the GBM stem cell state also comes from the most highly correlated individual gene with sensitivity, SOX9, which is a known regulator of stemness in GBM^58–61^. SOX9 is also an interactor of the NuRD complex which can impact histone acetylation^64^. Therefore, combination treatment sensitivity may be primarily related to the contribution that NuRD and SOX9 make to the GBM stem cell epigenome, an important area of future study.

Overall, this study builds on a robust mechanistic understanding of the interaction between the EYAs and PLK1, to provide mechanism-driven rationale for a novel combination therapy. We provide strong evidence that combined EYA and PLK1 inhibition is effective in cancer cell lines, cancer stem cell models, and tumour spheroids. Importantly, we lay the groundwork for biomarker-based therapeutic testing by demonstrating that the combination treatment specifically reduces the viability and growth of tumour models with overexpression of EYA family members and the NuRD complex. Finally, we demonstrate that the combination treatment specifically targets the GBM stem cell state, a finding of particular importance given the integral role of GBM stem cells in tumour formation and disease recurrence. As the first demonstration of combined PLK1 and EYA inhibitor efficacy, this work highlights the need for more potent second-generation EYA inhibitors that will help bring the strategy to clinical application.

## Methods

### Cell culture

#### Cancer cell lines

Cancer cell lines were either maintained in DMEM with 10% FCS in a sterile humidified incubator at 37_°_C with 10% CO_2_ and 20% O_2_ (SJSA, SAOS2, A172, H4, T98G, HEPG2, IMR32, SKNDZ, SKNAS, SW48, and CACO2), or RPMI with 10% FCS in a sterile humidified incubator at 37_°_C with 5% CO_2_ and 20% O_2_ (769P, Kelly, SKNFI, SNU61). Cells were passaged and harvested with trypsin. RNA expression levels for EYA1 and EYA4 were downloaded from https://depmap.org/portal/,22Q4.

#### GBM stem cells

GBM stem cell lines were acquired from QIMR Berghofer and grown in monolayers on Corning® Matrigel® cat #354234 coated flasks (1:100 dilution) and maintained in StemPro® NSC SFM cat #A1050901 (Thermo) as per manufacturer instructions and with supplemental epidermal growth factor (20 ng/ml) and fibroblast growth factor (10 ng/ml). Cells were passaged and harvested with accutase. Stem cell differentiation was performed by replacing growth factor containing media with media supplemented with 2% FCS for 48 hrs, and reseeding of cells in 2% FCS prior to further experimentation.

#### Generation of T98G FUCCI(CA)2 cell line

3-colour FUCCI T98G cells were generated using the tFucci(CA)2/pCSII-EF construct (RDB15446, provided by H. Miyoshi through RIKEN BRC^65^. Purified aliquots of third-generation lentiviral vector containing the construct (Vector and Genome Engineering Facility, Sydney, NSW, Australia) were transduced into T98G cells in a T-25 flask at a multiplicity of infection of 3, supplemented with 4ug/mL polybrene (Sigma-Merck) for 24 hours in DMEM. Cells were expanded for 5 days and sorted for mCHERRY or mVenus expression (Flow Cytometry Facility, Westmead Institute for Medical Research, Sydney, NSW, Australia). Sorted cells were seeded into a 24-well plate with 1mL media supplemented with Penicillin-Streptomycin (Gibco). The T98G-FUCCI cells were expanded for a further 7 days, and sorted once more to obtain a pure population of expressing cells. The purified T98G-FUCCI cells were passaged in media supplemented with Penicillin-Streptomycin for an additional 7 days, after which the Penicillin-Streptomycin was removed and cells cultured as normal. Fluorescence and corresponding cell cycle-dependant signalling was verified using live cell imaging.

### AlamarBlue viability assay and drug synergy

Cancer cell lines or PB1 stem or differentiated cells were seeded in 96 well plates at 15-20% confluence on day 1. On day 2, cell media was replaced with media containing drugs at the specified concentrations. DMSO concentration was matched across all treatments. Following 72 hr incubation with drugs, cell viability was measured using alamarBlue™ HS cell viability reagent (#A50101, Thermo Scientific) according to manufacturer’s instructions. Statistical comparisons between groups were made using two-sample t-tests to compare the mean viability at each dose of drug or combination. ZIP synergy scores were computed using the web-based synergyfinder tool: https://synergyfinder.org/#!/, or https://synergyfinder.fimm.fi/synergy/2024122302535356633/. Statistical comparison between mean ZIP synergy scores were performed using two-sample t-tests.

### GBM tumour spheroids

To grow GBM stem cells as tumour spheroids, cells were seeded at 1500 cells/well into round bottom 96 well plates in GBM stem cell media. Spheroids were grown in the Incucyte S2® live-cell analysis system and imaged every 8 hrs using the spheroid setting. After 125 hrs, media was replaced with media containing a final concentration of the following drugs: benzarone (7.5 µM), BI2536 (10 nM), volasertib (20 nM), combinations of drugs as indicated, or DMSO. The final concentration of DMSO was matched across all treatments. Media was replaced with fresh media with or without drugs depending on the timepoint every 48-72 hrs throughout the experiment. Five PB1 spheroids were included per treatment group at time 0, however a few spheroids were aspirated during media changes throughout the experiment and only spheroids that persisted for the duration of the experiment were included in the final analysis (at least 3 spheroids/treatment group). Mean spheroid area and standard deviations were plotted over time for each treatment group. Final spheroid area was compared between combination treated spheroids and all other treatment groups using ANOVA and Šídák’s multiple comparisons test. Sporadic missing measurements for individual spheroids occurred due to failure of the Incucyte autofocus function.

### Live cell proliferation experiments

#### Confluence

Cancer cell lines or GBM stem cell models were seeded into 96 well plates or matrigel-coated 24 well plates, respectively. On day 2, cells were treated with drugs at the indicated doses and imaged in an Incucyte zoom® or Incucyte S2® live-cell analysis systems at regular intervals for either 3 days (cancer cell lines), or for 4-5 days with a drug replacement following the first 2 days (GBM stem cell models). Cancer cell line mean confluence was quantified over time were plotted over time and the growth area under the curve was compared between treatment groups using ANOVA and Tukey’s multiple comparisons test. GBM stem cell confluence over time was used to compute the total relative growth for each treatment. Means are presented with standard errors and were compared using ANOVA and Tukey’s multiple comparisons test.

#### T98G FUCCI(CA)2 cells and Annexin V

T98G FUCCI(CA)2 cells or WT T98G cells were seeded into 96 well plates. On Day 2, the DMEM was replaced with FluoroBrite™ DMEM containing the dose response matrices and subsequently imaged. For Annexin V analysis, cells were simultaneously treated with Incucyte® Annexin V Green Dye (Catalog #4642) at 1:200 dilution according to manufacturer’s instructions. Cells were grown in the Incucyte S2® live-cell analysis system. Imaging and analysis were performed using the Incucyte® Cell-By-Cell Analysis Software Module to mask individual cells and classify them into different cell cycle stages or by the presence of the Annexin V dye signal. The mean percentage of cells in each cell cycle stage or positive for Annexin V were plotted with standard error of the mean. Additionally, heat maps corresponding to all doses within the dose response matrix were plotted for G2/M cells and Annexin V positive cells at both 16 hr and 72 hr timepoints.

### Immunoblotting

Western blot lysates were produced in RIPA buffer (Pierce #89900) supplemented with cOmplete Mini EDTA-free protease inhibitor cocktail (Roche) and 1 mM dithiothreitol at 4 °C. Lysates were cleared by centrifugation at 16,000 x g for 30 minutes at 4 °C. Proteins were separated using 3-8% or 7% Tris-Acetate, or 4-12% Bis-Tris precast mini gels (Life Technologies) and transferred to PVDF membranes at 70 V for 2 hrs (Immobilon P, Millipore). Ponceau S staining was used to verify even transfer (Sigma Aldrich). Membranes were optionally cut to detect multiple proteins. Blocking and antibody incubations were then performed with 5% non-fat milk in PBST. Bands were visualized with HRP-conjugated secondary antibodies (DAKO) followed by application of a chemiluminescent reagent (Thermo Scientific). Stripping with Restore^TM^ PLUS stripping buffer (Thermo Scientific) and reprobing was performed in some cases when the subsequent primary antibody was from a different species, however, blots were never stripped more than once. Western blot primary antibodies targeted EYA1, Protein Tech 22658 (1:1000), EYA2, Millipore HPA027024 (1:1000), EYA3, Protein Tech 21196 (1:1000), EYA4, Santa Cruz SC393111 (1:1000), Actin, Sigma a2066 (1:2000), PARP, Cell Signalling 9542 (1:1000), gamma-H2AX Millipore 05-636 (1:1000), Vinculin, Sigma V9131 (1:4000), p-S10 H3 Cell Signalling 9706 (1:1000), CD133, Cell Signalling 64326 (1:1000), and Sox9, Cell Signalling 82630 (1:1000).

Densitometry was performed in ImageJ on unaltered images. Values were statistically compared where indicated using two sample t-tests. Correlation analysis between Z-scores of individual or combinations of EYAs and sensitivity to the combination treatment (Fig. 4F) was performed in Microsoft excel using the correl function.

### CETSA assay

Following the indicated drug treatments, cells were trypsinized and diluted in PBS with cOmplete Mini EDTA-free protease inhibitor cocktail (Roche) in 9 PCR tubes per treatment. Tubes were heated to the indicated temperatures for 5 minutes using a thermocycler followed by incubation at room temperature for 3 minutes and snap freezing with liquid nitrogen. Lysates were prepared by thawing at room temperature, vortexing for 10 seconds, and snap freezing in liquid nitrogen for 5 cycles. After the final freeze-thaw cycle, samples were vortexed for 1 min each, and then lysates were cleared by centrifugation at 16,000 x g for 30 minutes at 4 °C. Cleared lysates were used for western blots, as described. Densitometry for each EYA is expressed relative to each treatment’s 37°C sample.

### Immunofluorescence experiments

Cells were grown on coverslips pre-treated with Alcian blue stain to promote adherence (Sigma) and treated with the designated drugs for 5.5 hrs. For p-S46 TCPT staining, cells were washed with PBS and then fixed using freshly prepared 4% paraformaldehyde (Sigma) for 10 minutes at RT followed by 2X PBS washes and then permeabilization with KCM buffer (120 mM KCl, 20 mM NaCl, 10 mM Tris pH 7.5, 0.1% Triton). Next, blocking was performed using antibody-dilution buffer (20 mM Tris–HCl, pH 7.5, 2% (w/v) BSA, 0.2% (v/v) fish gelatin, 150 mM NaCl, 0.1% (v/v) Triton X-100 and 0.1% (w/v) sodium azide) for 1 hr at RT. Cells were incubated with anti-p-S46 TCTP (Cell Signalling 5251) and anti-p-S10 H3 primary antibodies (Cell Signalling 9701) overnight at 4_°_C followed by 3 x 10 minute washes in PBS. Cells were then incubated with Alexa Fluor conjugated secondary antibodies (Thermo Scientific) at a 1:750 dilution for 1 hr at RT. Cells were again washed for 3 x 10 minutes in PBS followed by staining in DAPI solution for 20 minutes (Sigma). Coverslips were mounted on slides in ProLong^TM^ Gold antifade.

RAD51 staining and imaging was similar except that EdU was added to the cell media for 1 hr prior to fixation at a final concentration of 10 µM, and Click-It EdU staining was performed as per the manufacturer’s recommendations (Thermo Scientific) immediately following fixation. Additionally, permeabilization in KCM was performed prior to fixation to remove loosely attached mitotic cells. Finally, the primary antibody in this experiment was anti-RAD51 (Abcam 63801).

Images were acquired with a Zeiss Axio Imager microscope at 40X. Cell, nuclei, and foci masking, counting and intensity measurements were performed in Cellprofiler version 4 using custom image analysis pipelines with all settings kept consistent across treatments and biological reps for each experiment^66^. All intensity measurements were made on raw .tiff images.

Cell cycle plots were made from the combined data of the two biological replicates (intensity values are relative to the DMSO mean within each independent rep). Cell cycle gates were determined based on DAPI intensity and either p-S10 H3 intensity or EdU intensity. Gating values for p-S10 H3 and EdU were consistent across all treatments within each replicate, whereas DAPI intensity values were determined based on subjective assessment of the G1 midpoint in each treatment, and then fixed percentile ranges based on the G1 midpoint were used to determine the min/max values of G1 and G2 or G2/M cells. Intensity and foci count data were presented for individual cells within specific cell cycle phases. Means were presented with standard errors. Statistical comparisons were performed using ANOVA with Tukey’s multiple comparison test.

### Sister chromatid exchanges

Cells were cultured in fresh medium supplemented with 10 µM BrdU (Sigma-Aldrich) for 40 hrs depending on the mitotic index of the cell line. Cell cultures were treated with 20 ng/ml colcemid for the last 4 hr of incubation to accumulate mitotic cells. Cells were harvested by trypsinization and centrifugation and then incubated in hypotonic buffer for 10 min at 37 °C. Swollen cells were fixed by gradually adding 1 mL of fresh ice-cold fixative (methanol/acetic acid 3:1), mixing by inversion and incubating on ice for 5 min. Cells were then collected by centrifugation at 1500 × g for 8 min. 10 mL ice-cold fixative was added to resuspend the cells followed by 5 min incubation on ice and centrifugation at 1500 × g for 8 min. This fixing step was repeated another two times. Fixed cells were then resuspended in 500–1000 µL of ice-cold fixative, and dropped onto clean, dry, microscope slides (HDS Surefrost, 50–100 µL cell solution per slide). To drop chromosomes, a clean dry slide was held over a 75 °C water bath, and the cell solution was dropped from a pipette onto the slide, which was quickly flipped and held close to the surface of water bath for 5 s. Slides were left to dry for 2 to 3 days and then treated with 100 μg/ml DNase-free RNase A (Sigma) in 2 × SSC for 30 min at 37 °C, rinsed in PBS, and postfixed in 4% formaldehyde in PBS at room temperature for 10 min. Following a quick rinse in deionized water, slides were dehydrated in a graded ethanol series (70% for 3 min, 90% for 3 min, and 100% for 3 min) and allowed to airdry. Slides were then stained in 0.5 μg/mL Hoechst 33258 (Sigma–Aldrich) in 2 × SSC for 15 min at room temperature, rinsed in dH_2_O, and air-dried. Slides were then flooded with 200 μL 2 × SSC and exposed to long-wave (∼365 nm) UV light (Stratalinker 1800 UV irradiator; Agilent Technologies) for 45 min. The BrdU/BrdC-substituted DNA strands were then digested in 10 U/μL Exonuclease III solution (New England Biolabs) in the supplied buffer at 37 °C for 30 min. After a quick rinse in deionized water, slides were incubated with 50 ng/mL DAPI in PBS for 15 min, washed twice in PBST for 5 min, rinsed in deionized water, and airdried. Airdried slides were mounted in Prolong Gold Antifade (Invitrogen) and stored at 4 °C until microscope analysis. Slides were imaged by automation on the MetaSystems Metafer Scanning Platform (Carl Zeiss) microscope.

### Correlation, gene-ontology, and pathway analysis

Proteomic data from the Q-cell data repository corresponding to the 12 GBM stem cell models (https://www.qimrberghofer.edu.au/commercial-collaborations/partner-with-us/qcell/) was first pre-processed in Perseus, including imputation of missing values using a width of 0.4 and a downshift of 1.8. Correlation coefficients and associated p-values between individual proteins and GBM stem cell line sensitivity to the combination treatment were determined in Microsoft Excel. Gene-ontology, and pathway analysis was performed using https://string-db.org/ and the proteins with values/ranks function (correlation coefficients were used as values). Gene-ontology terms and pathways with enrichment scores of > 4 were visualized with https://string-db.org/. The 15 core NuRD component subunits (listed in EV 6A) were also correlated with combination sensitivity across other datasets in Fig. 6E from the following sources: https://www.qimrberghofer.edu.au/commercial-collaborations/partner-with-us/qcell/ (all Q-cell datasets), Gonçalves et al., 2022 (pan-cancer cell line proteomics),^67^ and https://depmap.org/portal/ (pan-cancer cell line RNA-Seq).

## Acknowledgements

Dr. Brett Stringer and Dr. Bryan Day are thanked for generously providing the 12 GBM stem cell “QCell” models. We also thank Dr. Tony Cesare for providing protocols for FUCCI cell generation.

---

**EV Figure 1:**
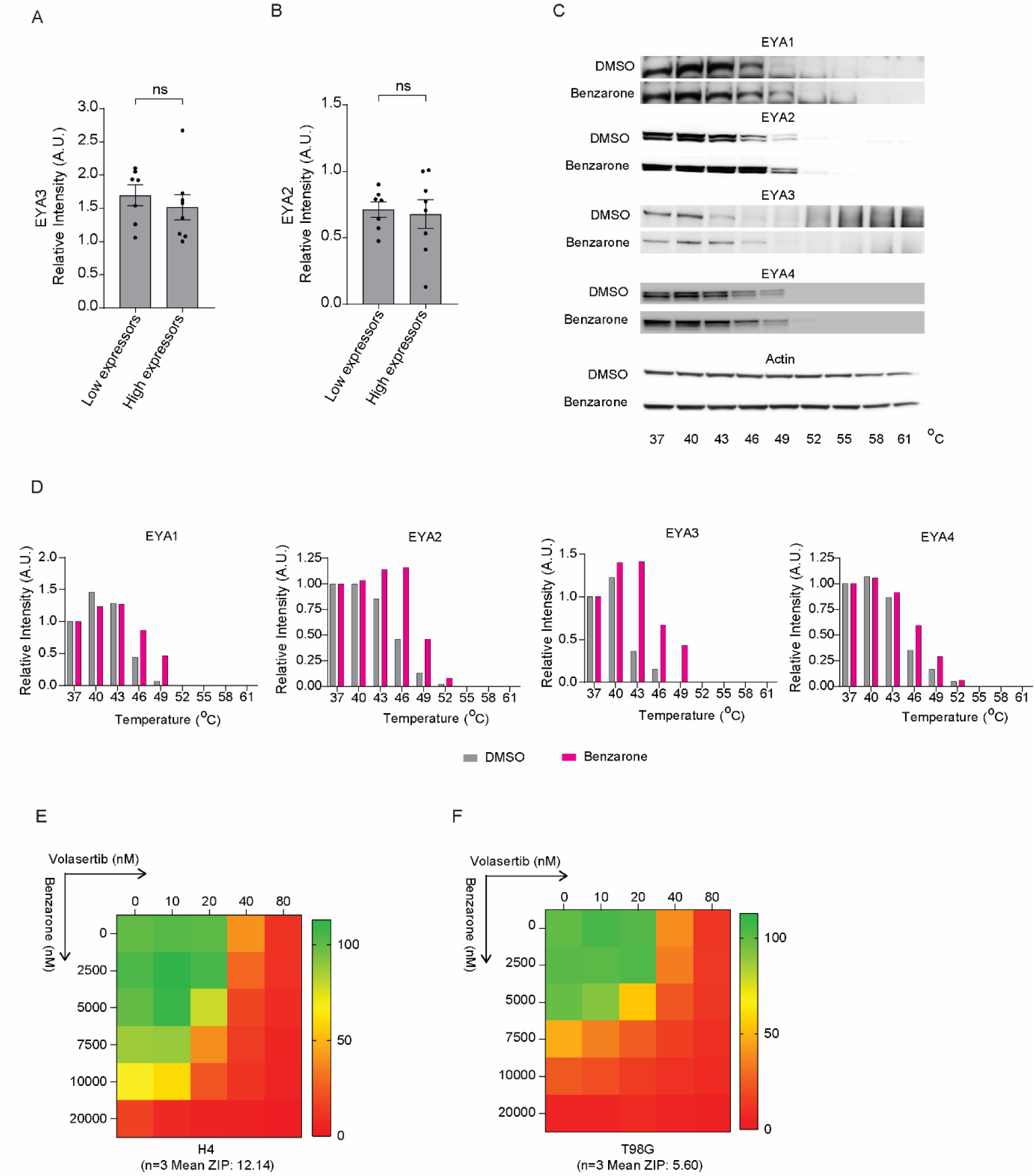
Supporting analysis of EYA expression levels, benzarone specificity, and an alternative PLK1 inhibitor-based combination. **(A-B)** Quantification of mean EYA3 or EYA2 protein levels relative to IMR32 via densitometry from Figure 1B (two-tailed *t* test, ns = non-significant). **(C)** IMR32 cell CETSA assay western blots of EYA1-4 following treatment with DMSO or Benzarone (10 µM, 2 hrs) and exposure of lysates to the indicated temperatures. **(D)** Densitometry quantitation of the CETSA assay. EYA1-4 band densitometry values are expressed relative to the respective value in the 37°C treatment. Greater values for each EYA at increasing temperatures in the benzarone treated samples compared to DMSO indicates target binding. **(E)** Relative viability across combination dose-response matrixes (Benzarone and Volasertib) measured by alamarBlue in H4 cells (n=3, mean ZIP synergy score is shown below) **(F)** Relative viability across combination dose-response matrixes (Benzarone and Volasertib) measured by alamarBlue in T98G cells (n=3, mean ZIP synergy score is shown below).

**EV Figure 2:**
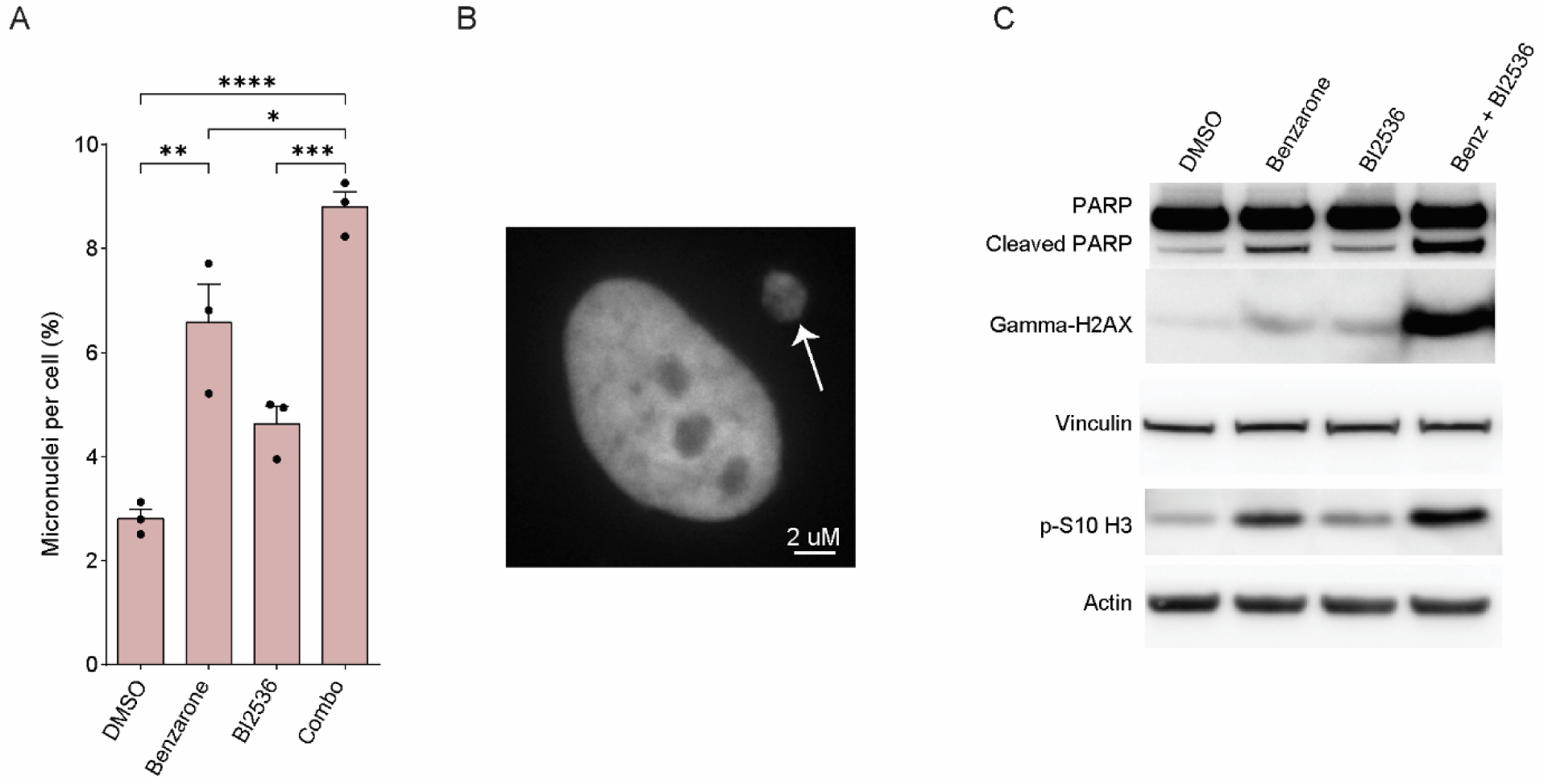
Combined EYA and PLK1 inhibition increases mitotic cell fraction, cell death, DNA damage, and micronuclei in H4 cells. **(A)** Percentage of cells with micronuclei in H4 cells following 24-hour treatment with DMSO, Benzarone (7.5 µM), Bi2536 (10 nM), or Benzarone + Bi2536 (one-way ANOVA with Tukey’s multiple comparison test, * p ≤ 0.05, ** p ≤ 0.01, *** p ≤ 0.001, **** p ≤ 0.0001). **(B)** Representative image of H4 cell with micronuclei (white arrow). **(C)** Western blots in H4 cells using the indicated antibodies following 24 hr treatment with DMSO, Benzarone (7.5 µM), Bi2536 (10 nM), or Benzarone + Bi2536.

**EV Figure 3:**
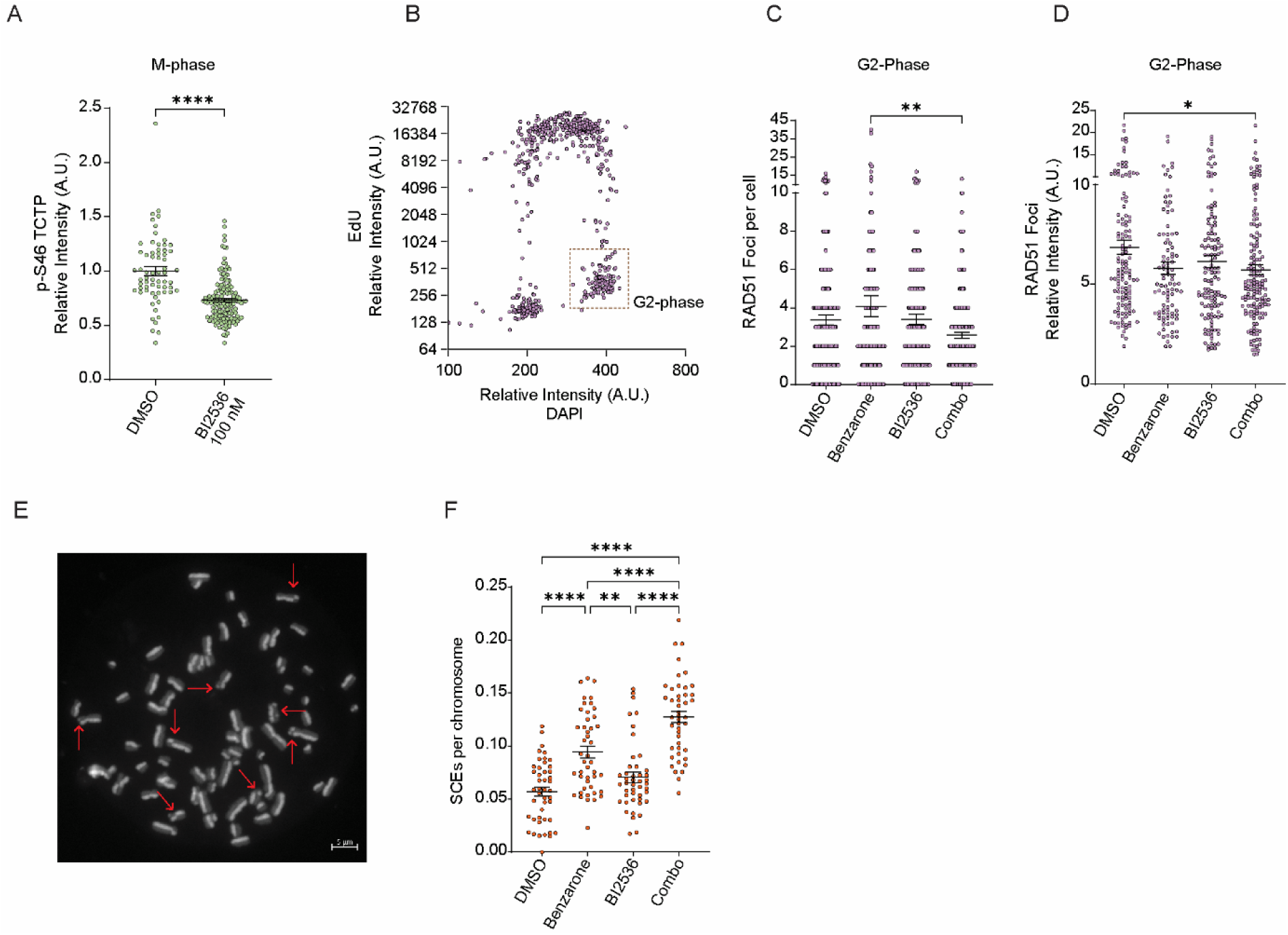
Combined EYA and PLK1 inhibition additively decreases RAD51 foci formation and intensity in G2-phase and increases sister chromatid exchanges. **(A)** Cellular quantitation of p-S46 TCTP intensity in M-phase following treatment with DMSO, Benzarone (7.5 µM), Bi2536 (10 nM), or Benzarone + Bi2536 for 5.5 hrs (two-tailed *t* test, **** p ≤ 0.0001). **(B)** Representative quantitative imaged based cytometry plot showing G2-phase cell cycle gate as determined by staining intensity of EdU and DAPI. **(C)** Quantitation of RAD51 foci number in G2-phase following treatment with DMSO, Benzarone (7.5 µM), Bi2536 (10 nM), or Benzarone + Bi2536 for 5.5 hrs (one-way ANOVA with Tukey’s multiple comparison test, ** p ≤ 0.01). **(D)** Quantitation of RAD51 foci intensity in G2-phase following treatment with DMSO, Benzarone (7.5 µM), Bi2536 (10 nM), or Benzarone + Bi2536 for 5.5 hrs (one-way ANOVA with Tukey’s multiple comparison test, * p ≤ 0.05). **(E)** Representative image of chromosome spread with SCEs indicated with red arrows. **(F)** Quantitation of SCEs per chromosome following treatment with DMSO, Benzarone (7.5 µM), Bi2536 (10 nM), or Benzarone + Bi2536 for 24 hrs (one-way ANOVA with Tukey’s multiple comparison test, ** p ≤ 0.01, **** p ≤ 0.0001).

**EV Figure 5:**
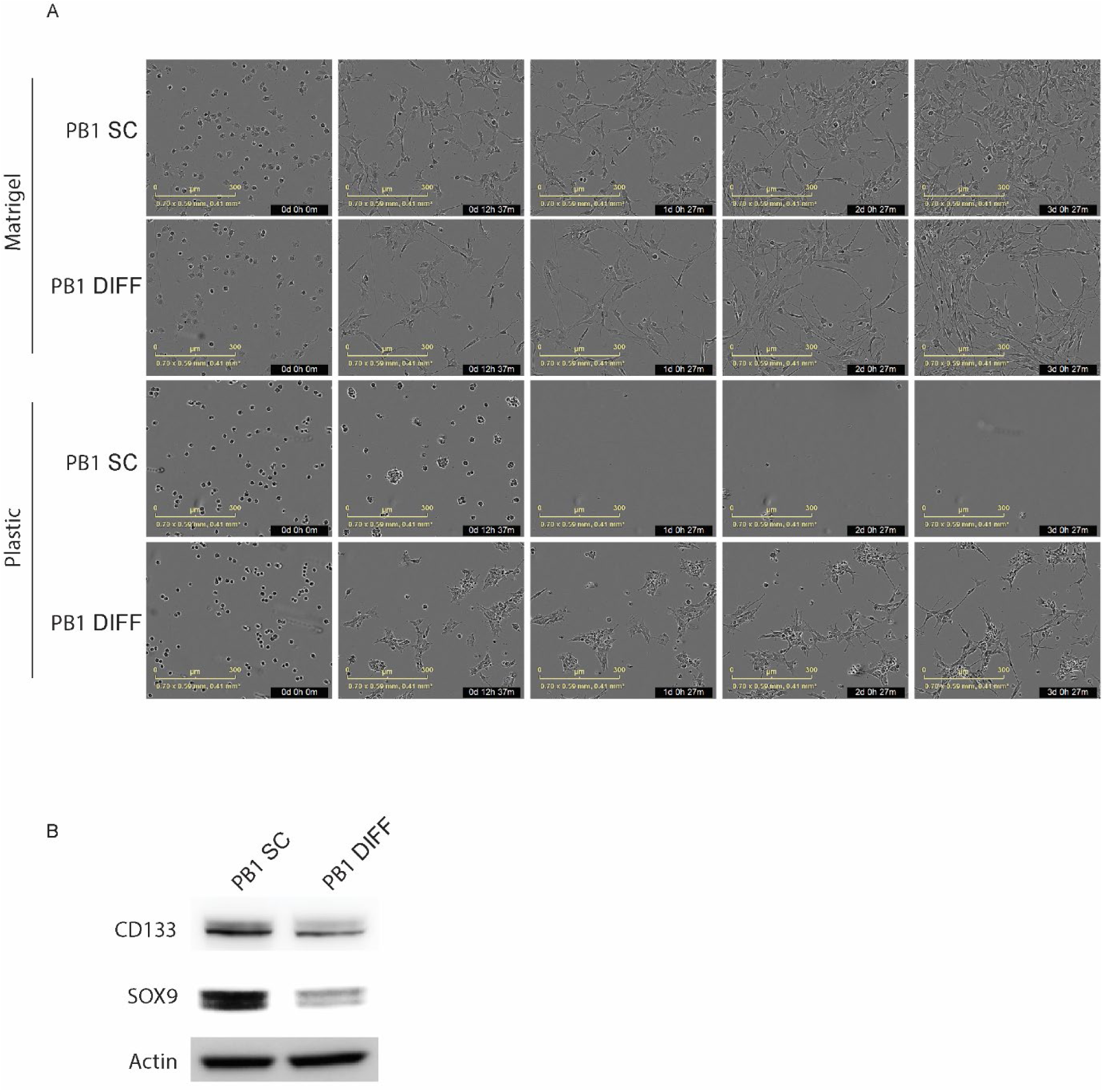
Validation of GBM stem cell differentiation. **(A)** Representative images over time of PB1 stem cell (SC) or differentiated cells (DIFF) grown on matrigel or standard tissue culture dishes (plastic) at the indicated timepoints. **(B)** Western blots of PB1 stem cell (SC) or differentiated cells (DIFF).

**EV Figure 6:**
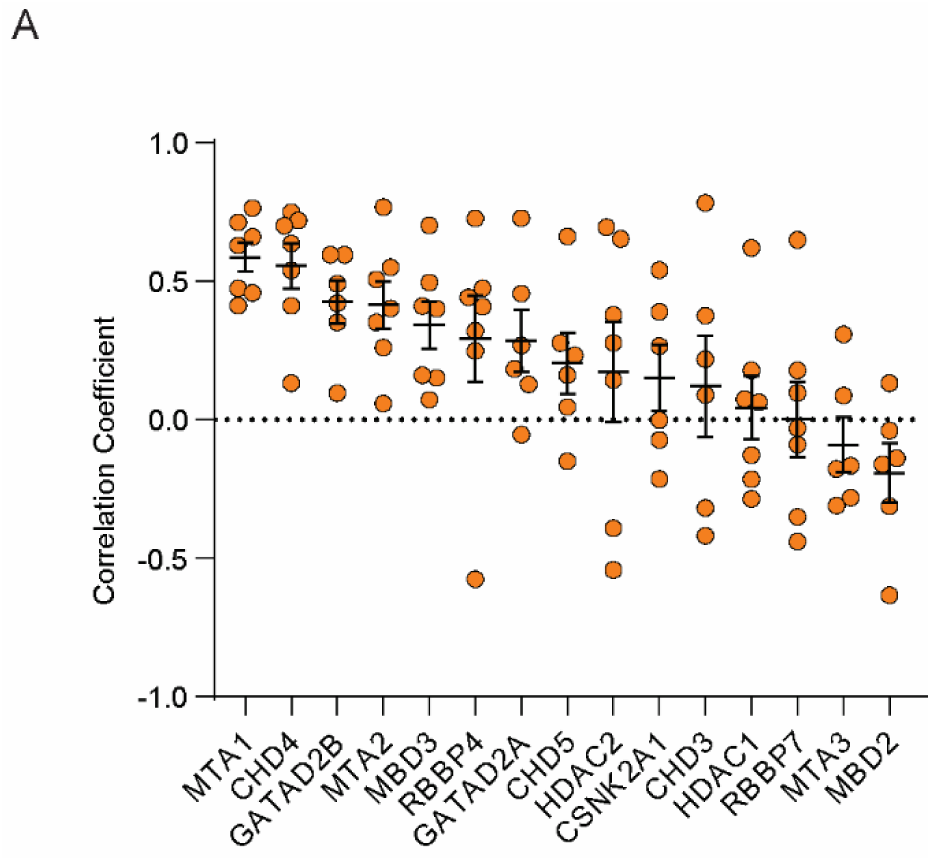
Individual NuRD component correlations across datasets. **(A)** Pearson correlation coefficients between combination viability loss and NuRD proteins across proteomic and transcriptomic datasets in GBM stem cell and pan cancer panels (each datapoint represents the correlation from a dataset listed in Fig. 6E).

